# The role of climate change education on individual lifetime carbon emissions

**DOI:** 10.1101/441170

**Authors:** Eugene C. Cordero, Diana Centeno, Anne Marie Todd

**Affiliations:** Department of Meteorology and Climate Science, San José State University, San José, California, United States of America; Department of Communication Studies, San José State University, San José, California, United States of America

## Abstract

Strategies to mitigate climate change often center on clean technologies such as electric vehicles and solar panels, while the mitigation potential of a quality educational experience is rarely discussed. In this paper, we investigate the long-term impact that an intensive one-year university course had on individual carbon emissions by surveying students at least five years after having taken the course. A majority of course graduates reported pro-environmental decisions (i.e., type of car to buy, food choices) that can be attributed to experiences gained in the course. Furthermore, our carbon footprint analysis demonstrates that for the average course graduate, these decisions reduced their individual carbon emissions by 2.86 tons of CO_2_ per year. Focus group interviews identify that course graduates have developed a strong personal connection to climate change solutions, and this is realized in their daily behaviors and through their professional careers. The paper discusses in more detail the specific components of the course that are believed to be most impactful, and it shares preliminary outcomes from similar curriculum designs that are being used with K-12 students. Our analysis also demonstrates that if similar education programs were applied at scale, the potential reductions in carbon emissions would be of similar magnitude to other large-scale mitigation strategies such as rooftop solar or electric vehicles.

## 1. Introduction

In 1992, the United Nations Framework Convention on Climate Change stated, “Education is an essential element for mounting an adequate global response to climate change” (UNFCCC 1992). Few would argue against the importance of education in providing an informed response to environmental problems. Solutions to climate change tend to focus on mitigation and adaptation measures, and successful implementation of either strategy requires an informed and educated citizenry. Yet despite the notion of education’s importance in responding to climate change, education is rarely mentioned in discussions of today’s major climate solution strategies. One reason that education programs do not often feature in discussions about climate change mitigation is that education programs typically have insufficient data to verify reductions in carbon emissions. This is in contrast to technologies such as renewable energy generation and the electrification of automobiles that can demonstrate reductions in carbon emissions using data. Should education be shown to be an effective tool to reducing emissions via changes in behavior and attitudes, it would seem likely that funding and interest in such methods would become more widespread and well supported.

Education has been found to be one method for promoting behavior change, but only under certain circumstances (e.g., Kollmuss and Agyeman 2002; Lyons et al. 2006). The environmental education literature offers some insight into the connections between education and behavior change, and it also provides guidance on how to encourage pro-environmental behavior (D’Amato and Krasny 2011; Craig and Allen 2015). The notion that knowledge leads to awareness and then to action has been countered with studies that document that knowledge and skills are not enough to change behavior (e.g., Bray and Cridge 2013). The literature suggests that more personal factors such as a deep connection to nature, personal relevance to the issue and personal agency towards action are important factors that contribute to successful behavior change programs (e.g., Hungerford and Volk 1992, Kollmuss and Agyeman 2002, Bamberg and Möser 2007). Even among successful programs, the question of how long the intended behavior is sustained can vary depending on the type of intervention, with longer and more sustained engagements tending to have more long-lasting impacts (Orams 1997). This previous research informs educational research programs towards designs that not only focus on information but also promote the personal qualities that can support sustained action.

The purpose of this paper is to evaluate the impact of an intensive university climate change course on individual long-term carbon emissions. The design of the course is described including aspects of the course that were intended to help students develop a deep connection with climate change and climate solutions. Five years of graduates from the course were surveyed at least five years after they took the course. The results of both survey data and focus group interviews provide an indication of the long-term impact of the course, and they contribute to our understanding of the potential role that education can play in long-term behaviors and attitudes. We then quantify the reductions in annual carbon emissions resulting from graduates’ pro-environmental behavior, and we compare the reductions achieved through this education program with other climate change mitigation measures. A discussion of how this type of educational approach may be used within the K-12 education system is also provided.

## 2. Methods

### 2.1. University Course and Students

In fall 2007, a new course was offered at San José State University (SJSU) that satisfied all three subject areas of the upper division general education (GE) requirements, plus the campus writing requirement. The course, COMM/ENVS/GEOL/HUM/METR 168 & 168W: Global Climate Change I & II (hereafter referred to as COMM 168), is taught over an academic year, with six credit hours in the fall semester, and three credit hours in the following spring semester. The course is taught in a standard lecture theatre and has about 100 students enrolled per year. The course uses a number of design approaches to impact students in ways that maximize effects on students’ personal and professional lives. This course has been taught every year since 2007 and continues to be popular with students.

The course evaluated in this research was taught by three faculty members from different departments with expertise in the core themes of climate science, climate mitigation and environmental communication. Although different professors taught the course during the five-year study period, the syllabus was consistent through the period.

The subject population consists of approximately 500 matriculated students from COMM 168 in the academic years between 2007 and 2012 and who received GE credit for the course (receiving a grade of “C” or better). The subjects are all now SJSU graduates and at least five years have passed since they took the COMM 168 course. Students were recruited through email via SJSU’s alumni database after campus Institutional Review Board approval was secured. The majority of students enrolled came from colleges outside of the sciences as shown in Table 1.

**Table 1.** The distribution of colleges from the reported major for each of the participants.

| **College** | **% of students reporting** |
| --- | --- |
| College of Social Sciences | 46% |
| College of Business | 18% |
| College of Humanities and the Arts | 18% |
| College of Applied Sciences and Arts | 11% |
| College of Science | 7% |

### 2.2. Survey and Focus Groups

An 18-item survey instrument (provided in Appendix A was developed to study participants’ beliefs about climate change and whether their own personal actions to mitigate climate change could be attributed to motivating factors associated with taking the COMM 168 course. Of the more than 500 students who took the course, 104 students from the five different course iterations between 2007 and 2012 completed the survey. The categories of questions focus on participants’ a) attitudes and beliefs about global warming and whether they perceive it to affect them personally, and b) whether any of the participants’ current pro-environmental behaviors can be attributed to taking the COMM 168 course. A follow-up semi-structured focus group interview was conducted with a small sample of participants to better understand survey responses.

### 2.3. Estimating Carbon Emission Reductions

Once responses to the survey questions were obtained, the potential carbon reductions from the decisions made by participants can be estimated. Details of the procedure used are provided in Appendix B, but we briefly describe the method here. We use an online personal carbon footprint calculator ((http://coolclimate.berkeley.edu/calculator)) to estimate how a particular action attributed to taking COMM 168 would impact individual annual carbon emissions. We start by calculating the annual carbon emissions for an average person in California. Then, based on the response to a particular question (e.g., participant attributed their current purchasing of renewable energy from their utility to the COMM 168 class), we use the calculator to determine the reduction in annual carbon emissions due to that particular action (e.g., participant reduced emissions by 1.38 tons/year by purchasing renewable energy from their utility). This procedure is repeated for each of the actions identified in the survey.

## 3. Results

### 3.1. Survey

The first part of the survey focused on participants’ attitudes and beliefs about global warming. A large majority of participants (83%) agreed with the statement, “Most scientists think global warming is happening.”, and most participants (84%) also felt that global warming would affect their lives “a great deal” or “a moderate amount.” This is notable since the general public often discounts the impacts that global warming will have on them personally (Semenza et al. 2008; Buys et al. 2012). Most participants (84%) also strongly agreed or agreed with the statement, “I have personally experienced the effects of global warming.”, and when asked about how global warming will affect future generations, 91% said “a great deal”.

The second group of questions asked about personal actions to reduce climate change and whether the COMM 168 course had any effect on those actions. The general areas of climate action included waste reduction, home energy conservation, transportation and food choices. Each question asked participants to reflect on how participation in COMM 168 may have affected their actions today in those areas.

A summary of the results for the different categories is provided in Figure 1. In the waste and home energy conservation categories, a large percentage of participants described engaging in some actions to reduce waste or reduce energy use in their home that they attribute to taking the COMM 168 class. This included recycling more often (95%), giving away or donating products so they can be reused (75%), buying products that have less packaging (64%), changing to more energy efficient light bulbs (86%) and purchasing energy saving appliances (59%). Fewer participants reported actions such as composting food scraps (48%), purchasing renewable energy from their utility (18%) and installing solar panels (4%).

**Figure 1.**
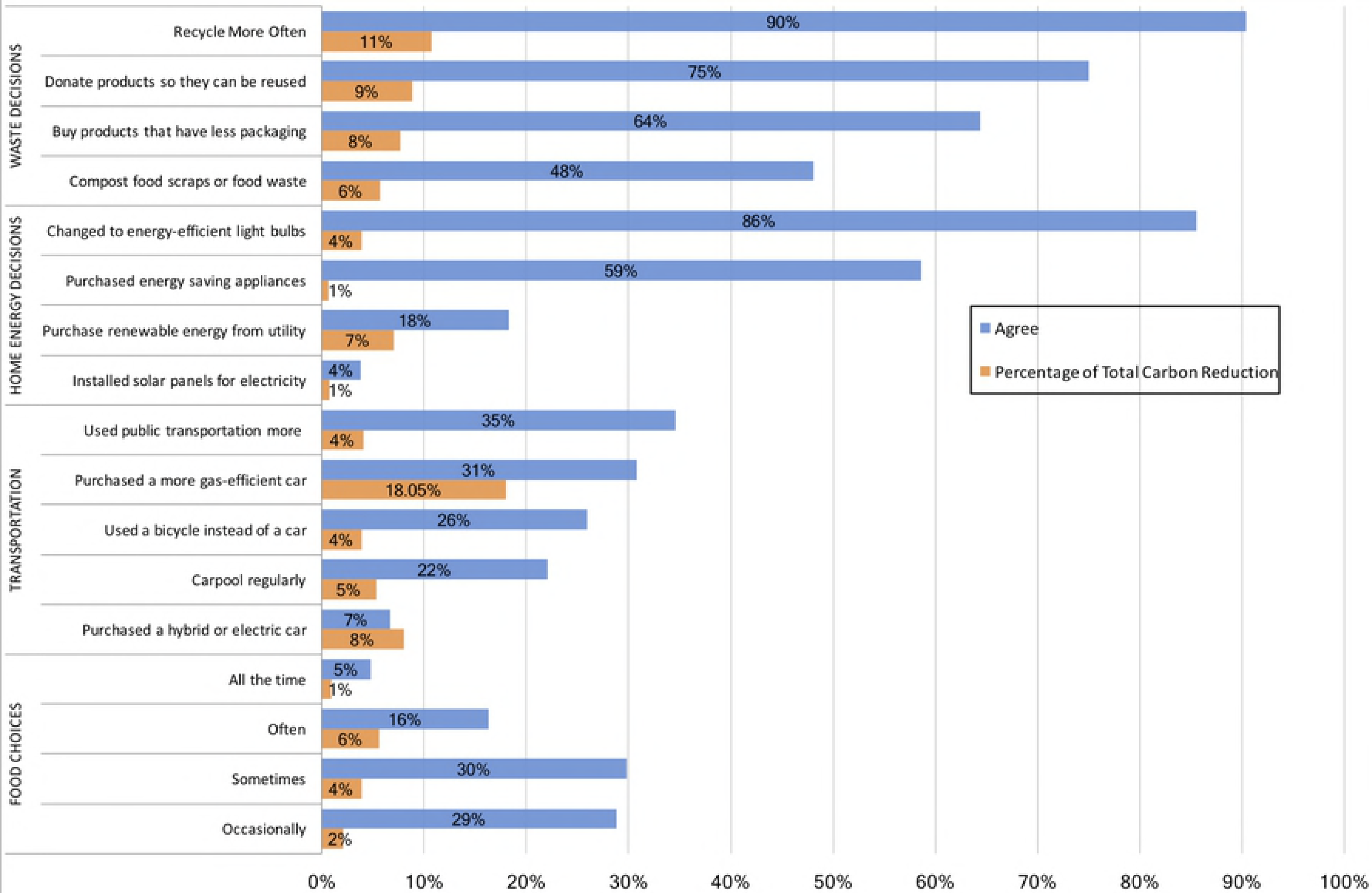
Survey results for questions related to how often participants make choices to reduce carbon emissions as a result of the COMM 168 class. The blue bars represent the percentage of students who agreed with the survey response, while the orange bars represent the relative impact in carbon emissions in percent relative to the total reductions.

In the transportation category, about 25% of participants reported some behavior to reduce emissions that is attributed to the COMM 168 class. This included using public transportation more (35%), using a bicycle for transportation (26%) and carpooling regularly (22%). And in the food choices category, most participants (80%) reported that at least occasionally they made food choices based on reducing carbon emissions.

The survey responses reported here suggest that participant behavior was influenced by the COMM 168 course in ways that continue to impact daily life. The types of actions studied here can be divided into two groups: one-time actions and recurring actions. For example, the purchase of an energy efficient light bulb or automobile is a one-time action, and these decisions will shape energy use for years into the future. In contrast, recurring actions such as recycling or food choices are made every day, and thus require more consistent engagement or behavior response. In reality, pro-environmental behavior includes both types of actions, and their impacts on carbon emissions can vary depending on the type of action and whether recurring actions become part of an individual’s lifestyle. Given the number of years that elapsed between the course and the survey, the survey provides a glimpse into behaviors that have likely become habitual. In the waste and food categories, some recurring actions were noted by most participants. Although recycling may be viewed as a fairly common action in many Californian communities, food choices and the connection with carbon emissions is not as widely known by the general public (e.g., Cordero et al. 2008; Wynes and Nicholas 2017). Given that 80% of participants reported some changes to their food choices, it appears that the course did have an impact on decision-making in this category even years after the class.

### 3.2. Estimated carbon emissions

Using the survey responses about the actions that participants took, we then calculate the reductions in carbon emissions for all participants. Figure 1 also shows the contribution of each of the survey questions to the total reductions in carbon emissions. While changes to behavior around reducing waste and energy conservation at home were the most common actions taken, the largest reduction in participant-averaged carbon emissions came through transportation decisions. For example, while only 31% of participants reported purchasing a more gas-efficient car, this single action accounted for 18% of all carbon emission reductions observed. In contrast, while over 90% of participants reported that they recycle more often, the combined reduction in carbon emissions only accounted for 11% of the total reductions.

As shown in Figure 2, the average reduction in carbon emissions based on the participant survey responses is 3.54 tons of CO_2_/year, with most participants between 2 and 5 tons of CO_2_/year. About 5% of students reported almost no change (0-1 ton of CO_2_/year), and about 10% reported between 6 and 8 tons of CO_2_/year. Of the four primary categories of carbon emission reductions, changes in transportation were responsible for 40% of the total carbon emission reductions, while waste reduction, food choices and home energy contributed respectively 33%, 13% and 12% of the achieved total carbon emission reductions.

**Figure 2.**
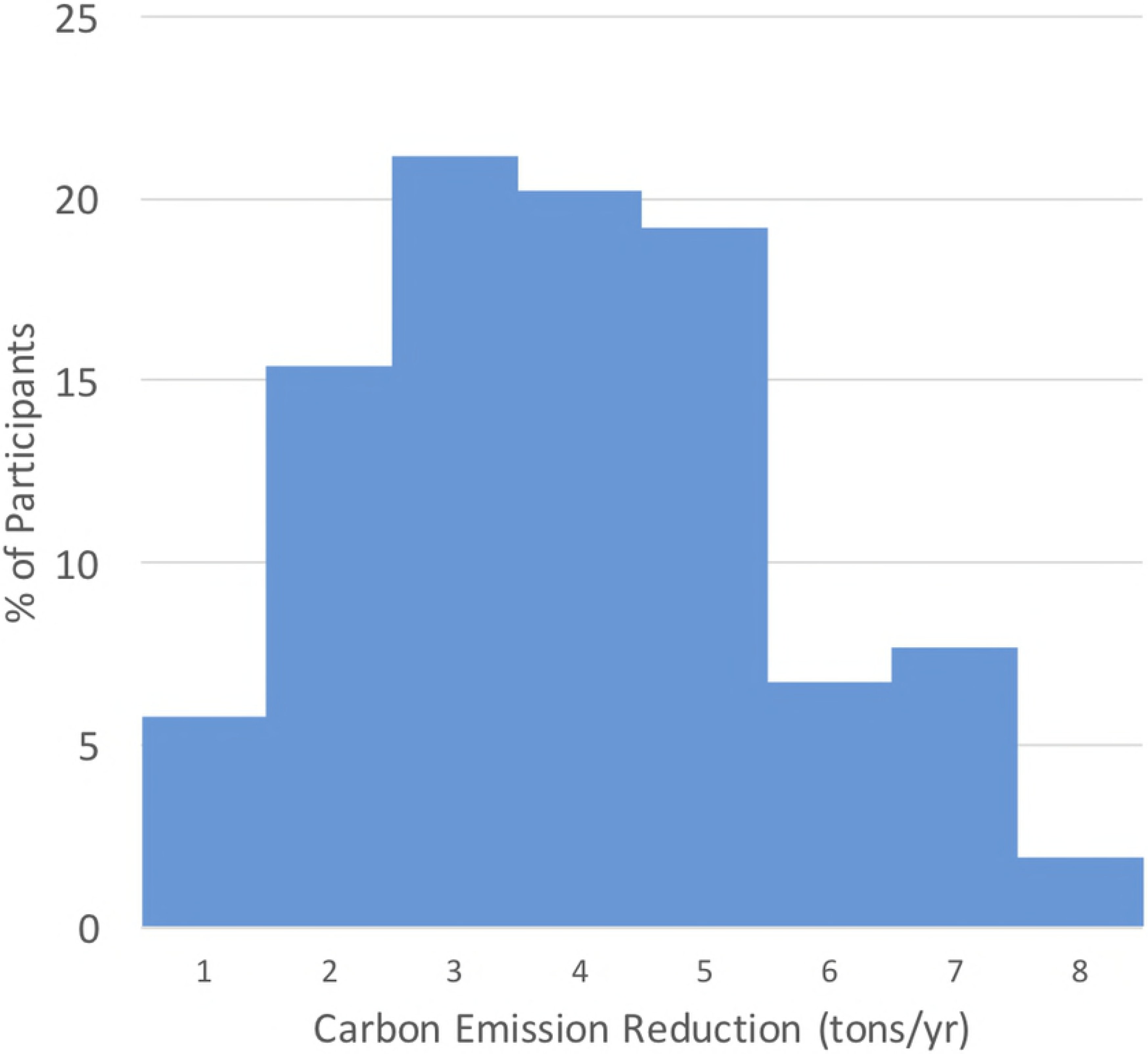
The distribution of carbon emission reductions (tons/year) for participants (n=104) as a result of the COMM 168 class.

### 3.3. Understanding how personal relevance and carbon emissions are related

Given that one of the goals of the course is to help students develop a personal connection between global warming and their lives, we explore the connections between participant beliefs and total carbon emission reductions through analysis of grouped data. In Figure 3, we show the relationship between individual carbon emission reductions with personal beliefs about how global warming will influence them or future generations. We find that participants who believe that global warming will harm them personally, or will harm future generations, have larger reductions in carbon emissions compared to participants who don’t believe there will be a strong impact on them or future generations. Thus it appears that in most cases, participants were at some level influenced by how they perceived the impact of global warming on their own wellbeing, or the well-being of future generations, when making personal decisions related to the environment.

**Figure 3.**
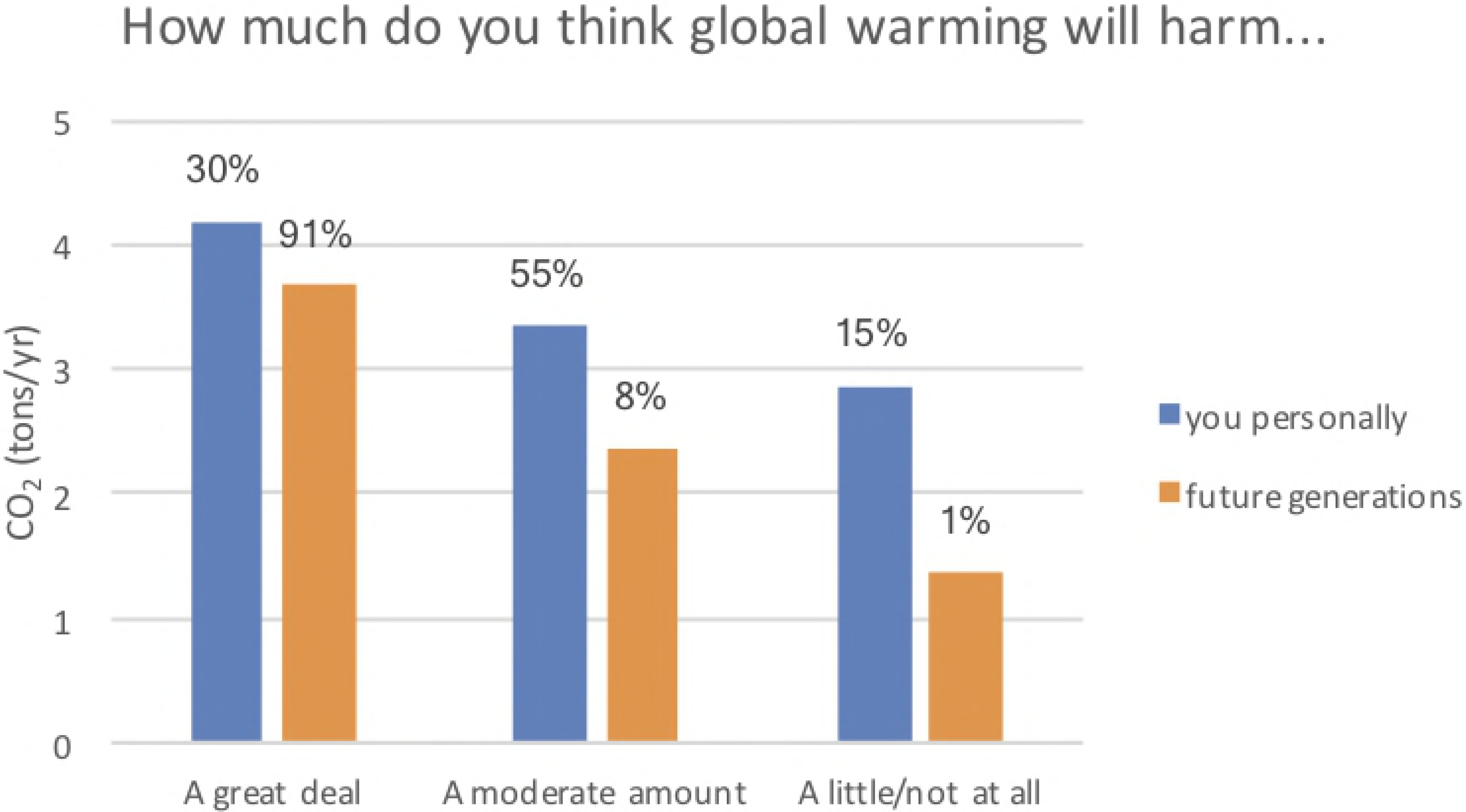
The reductions in carbon emissions (in tons/year) for groups based on their responses to the two questions about how global warming will affect them personally or future generations. The percentage of the total responses for that question is also given above each bar.

Further, of the participants who agreed (strongly agreed or agreed) to the statement, “I have personally experienced the effects of global warming,” their reductions in carbon emissions were 3.7 tons of CO_2_/year, while for the participants who didn’t agree with that statement (disagreed, strongly disagreed or neutral), their reductions were only 2.9 tons of CO_2_/year (see Figure 4).

**Figure 4.**
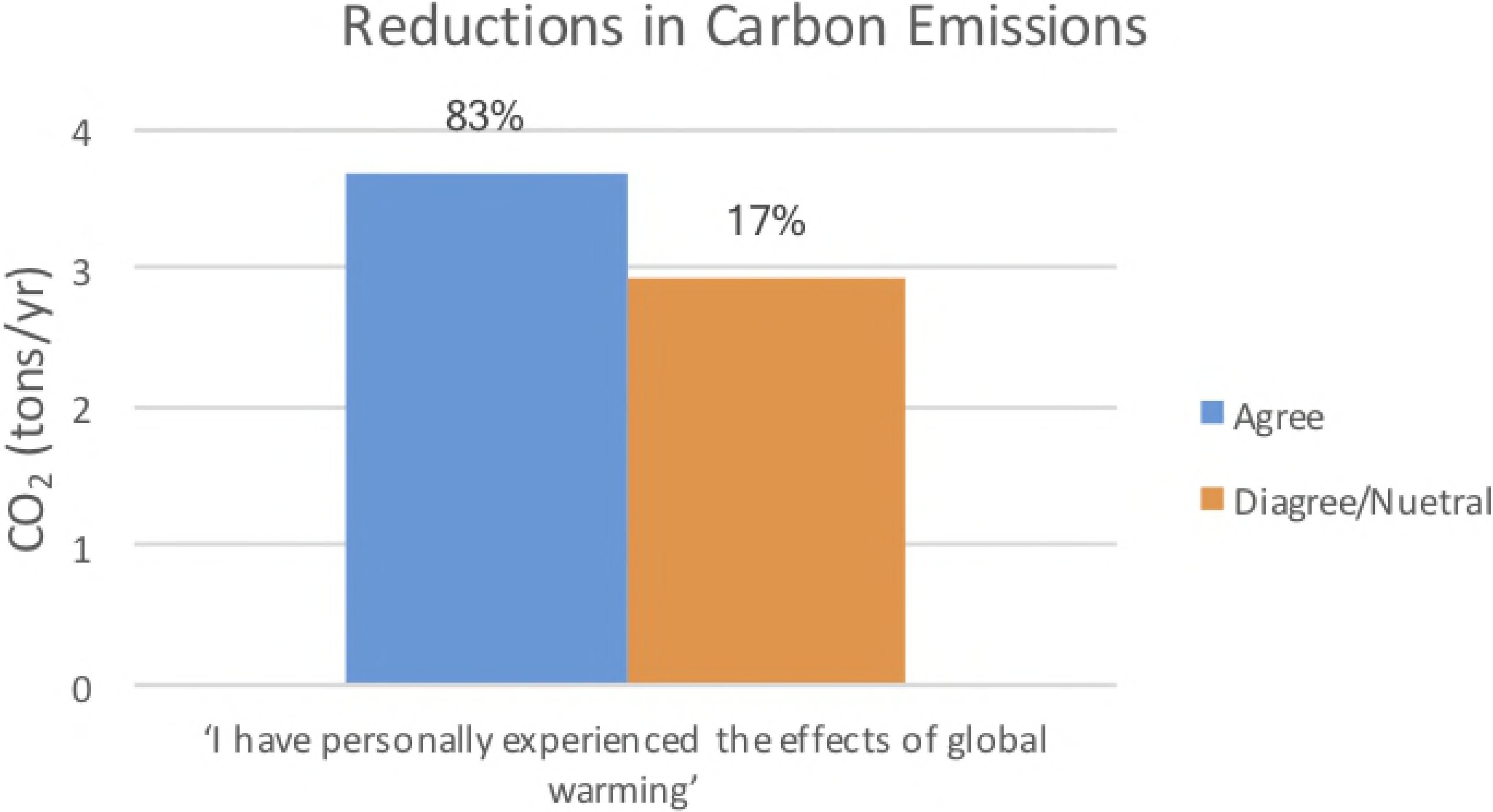
Relationship between statement response and total carbon emission reductions. For the first two statements, agree means strongly agree + agree, while disagree means strongly disagree + disagree + neutral.

### 3.4. Focus Groups

After evaluating the survey responses and noting themes in the utility of the course and personal climate change mitigation strategies, we followed up with focus group interviews to gain more in-depth understanding of the enduring influence of the course on students’ personal and professional lives. Including a qualitative approach such as focus group interviews can complement the survey analysis and ultimately enhance the quality of the resulting analysis (Wolff et al. 1993).

Focus group participants were recruited from 100+ survey respondents. We conducted two focus groups with a total of five participants. The focus groups were conducted in a classroom at San José State University. Participants were asked a series of open-ended questions about the course and its impact on their current lives. Responses from focus group participants converged around two themes: the importance of daily decisions to mitigate their climate change impact and the importance of engaging their community through climate change communication.

#### 3.4.1 Impact on Daily Decisions

A hallmark of conversations with graduates from the course was the consideration of climate in daily decisions. Fundamentally, focus group participants recognized the pervasiveness of climate change. As Tara, a focus group participant, noted, “Almost every activity we choose can affect [climate change] in some way. Whether we choose to take the bus or drive to work or whether we choose to buy food that’s grown on land that was cleared from rainforests…. Since it is in every aspect of our life pretty much, that automatically makes it relevant to all those different aspects.” Other participants agreed and described daily actions that centered on transportation, waste and food choices. Melissa noted, “I think about it all the time…. Definitely how I think about and go about my days, making decisions, even just from using plastic.” These responses exemplify a common theme in the group—the knowledge of climate change gained in the course prompted them to think about the impact of their actions.

The focus group participants noted that they go out of their way to take action because they feel as if they are making a difference. Billy noted, “When everyone does something to mitigate climate change, it will have a huge impact.” Tara concurred, “Almost everything I do can affect the climate somehow. If you start realizing how everything ties together, then pretty much everything you do, every choice you make can affect it in some way.” She continued, “I think every small step does make a difference…. One little step at a time, it all adds up. I’d like to think we’re making a difference. I feel like I am when I contribute a little bit.” Participants suggested that the interdisciplinary focus of the course allowed them to see the connections between their actions and broader climate forcings.

Participant comments demonstrate that environmental actions are not just because of sacrifice but that people feel good about taking action. Lolitta explained, “I feel good about how I’m living my life, and I’m excited by all the changes that I’m making, and I will continue making these changes because they make me feel great.” Billy noted proudly that he acts because “it’s like a moral obligation.” Ultimately, participants concurred that daily actions matter, and they cited this belief as the reason they continue to take actions. Their comments suggest they are empowered to act because they see themselves as part of the solution.

#### 3.4.2 Community Engagement through Communication

Overall, focus group participants noted that this course helped them develop experience communicating with other people about their actions and why they are taking them. Participants cited the community action class project as a key element in their understanding of the impact of community engagement. Examples of community actions projects include developing community gardens in a nearby apartment complex, giving presentations about climate solutions to schools, and implementing composting programs in nearby businesses. Billy described the lasting impact of “the hands-on approach” of the project: “Those experiences, I think for me, I carry those longer than / more than being in the classroom…. Being with people, doing something that’s going to translate into what I have to do work-wise in the future, [the project was a] translatable experience to the workforce.” Melissa described the lasting impression of the project as a crucial aspect to seeing the impact of action: “It actually bridged the gap between the course and what the community itself is doing”.

Participants noted the impact of the class on life beyond the home. One focus group participant, Elaine, is a manager at Walmart, and she credited the course for her “awareness in an industry with high consumption…. It’s so interesting how much I’ve been able to use just from this course.” She noted her focus as a manager is how to “reduce your inventory, reduce the waste, sell what you need to.” Elaine views Walmart’s waste issues as a management problem: “I see the huge amounts that they’re throwing away because they’re not managing their business correctly, because they’re not managing their production versus what they need, what they don’t. So that’s one of the things that I work on.” Elaine’s comments exemplify how many graduates of COMM 168 viewed the importance of taking action.

Billy noted the course explained how to make “big issues” like climate change “resonate with your audience…. That’s what I do now.” He explains that in his job at Pacific Gas and Electric, one of his roles is communicating about water issues, “that’s my biggest takeaway from this class: messaging. [I now understand] the importance of communicating about climate change in a way where people who don’t have a background in that subject can understand.” Other participants concurred that the course made them experts in climate change, and they now have to think about how to communicate with people who don’t have such extensive knowledge.

Participants also noted the importance of communicating with others about the actions they take. Tara noted: “It doesn’t really help unless you try to bring it out there. If I only ever walk places, no one will ever know unless I try to let them know why I walk places…. If you’re going to make a point by breaking the rules, you first have to know the rules because otherwise it doesn’t mean anything. If I want to rebel by not using a car, I first have to know that everyone thinks using a car is a normal thing to do.” Participants agreed that talking about their own actions helped in discussing climate change issues with others.

The community action project was a key part of the course in giving students experience outside of class in creating change. It also gave them some agency over this issue. Participants described their attempts to make a difference, both in their personal and professional lives. Participants noted the community action project allowed them to see the importance of communication in the design of their projects. The focus group responses suggest that interdisciplinary education including aspects of communication can give students the skills and experience necessary to create change in their own communities.

### 3.5. Educational approach

We have identified a number of key design elements that stood out as critical to the success of the education program we developed and that have sustained student engagement over many years. These include a) connecting climate science to students’ lives, b) providing students with experience creating change in a community of their choice and c) creating a culture devoted to stewardship and action. We found that these three key elements of the course helped students to connect with the subject in ways that extended into their personal and professional lives, and persisted over time to an extent that we have not seen in other similar classes. These elements were not isolated from each other, or from other important elements of the class, including a solid focus on climate science, climate mitigation and environmental communication. These factors are in line with the models suggested by other researchers, including personal relevance and empowerment (Bray and Cridge 2013). We now review each of these elements in more detail to provide insights into how these ideas may be applied to other educational settings.

#### 3.5.1 Connecting science to students’ lives

There were various activities in the class designed to help connect climate change with students’ lives. One project asked students to reflect on how climate change would affect their personal and professional lives. Another project had students track their personal energy use, and then implement a plan to reduce their energy use in their home using data from their home smart meters. These elements appeared to have some lasting impact, as various focus group participants reflected on how the course materials affected their personal and professional lives. In addition to the actions that were identified in the survey data, open-ended feedback also revealed that the course affected other major decisions, such as where to live and how many children to have. In fact, two of the participants mentioned their decisions to not have children or to adopt a child were influenced by the course. This implies that at least for some of the students, the course content and the implications of climate change affected their personal lives deeply. It appears that some of the high-impact actions identified by Wynes and Nicholas (2017) did resonate with the COMM 168 students.

#### 3.5.2 Creating change in a community of their choice

Another design element of the course was to provide students with real world experience creating and implementing an action plan to reduce carbon emissions. The Community Action Project (CAP) was the culminating experience during the second semester where student teams competed to develop the most impactful community-based project. The groups presented their results at the end of the semester, and were judged by an outside panel of experts. The goal of the CAP was to give students real-world experience developing solutions to climate change. It was our intention that through this experience, students would not only better understand some of the challenges associated with creating change but also gain confidence that change can happen through well-designed efforts. Ramsey (1993) found that using issue instruction and action training was an effective way to promote pro-environmental behavior. And MacFall (2012) found that students were deeply affected by their service-learning course even years after the experience. The COMM 168 class was focused around the year-long CAP and feedback from the focus groups shared how impactful the project was for some of the students, as a majority of the focus group participants mentioned the CAP as the most memorable part of the course. It is our conclusion that the CAP was an effective culminating experience, and today the class continues to use this activity to help students take their learning into practice.

#### 3.5.3 Creating a culture devoted to stewardship and action

The third aspect of the class that we feel was important to creating impactful and lasting change was the social norms that were established during the year-long class. Although we didn’t create any design elements at the start of the class to promote this, we do feel this was an important part of the course success. Among the emerging theories of behavior change, community-based social marketing (Mckenzie-Mohr, 2000; Heimlich and Ardoin 2008) and the strong impact of social norms have been shown to be effective measures for stimulating behavior change.

Creating a culture for a class is not easy, but there are effective teaching strategies that can help influence class culture, and we outline these here.

1) Group discussions with different students: We did a lot of group work in class. With 80-120 students per class, we took special efforts to mix students for their group work. This helps students work with new students and not develop their own group that they work with exclusively. We felt this helped students see multiple views across the class, and if an emerging interest and dedication to climate change arose through the class, it could spread.

2) Committed faculty: The faculty who taught this course were all deeply committed to climate change solutions, and they shared their own journey towards reducing carbon emissions. Through this one-year class, students got to know their professors well, and each professor engaged in some level of social action on their own. Role models are important in creating social change, and we suggest that having professors committed to environmental solutions was also a factor in creating social norms. For example, one of the focus group participants mentioned that as a student she used to drive an SUV, but she now drives a Prius. One of the professors said, “I didn’t know you drove an SUV when you took this class,” and she said, “I know you didn’t know because I didn’t want you to know.” This is an example of the social norms that were established in the class that may have extended to students’ lives outside of school and over time.

## 4. Potential Role of Education on Carbon Emission Reductions

Given the reductions in carbon emissions calculated in Section 3.2 (and shown in Figure 2), we now explore the potential role of education as a climate change mitigation strategy. We start by estimating the participant reductions in carbon emissions compared to a control group. The control group is created by using recent California’s per capita carbon emissions data as estimated by the California Air Resources Board (CARB 2017). Their data show that by 2014, per capita carbon emissions for the average Californian declined by 0.68 tons/year compared to 2009, the midpoint when students had graduated from SJSU. By contrast, the participants in COMM 168 reduced their per capita emissions by 3.54 tons/year. Thus, if we subtract the emission reductions for the average Californian (0.68) from our participants (3.54), we find that the net reduction above the average citizen is 2.86 tons/year (3.54-0.68 = 2.86).

We now use the net reduction in carbon emissions observed for graduates of COMM 168 to compare the potential role of education as a climate change mitigation strategy with other climate change mitigation strategies. For this comparison, we employ the methodology outlined in Project Drawdown (http://www.drawdown.org), where 80 different technologies or strategies are evaluated based on the potential to cumulatively reduce carbon emissions by 2050 (Hawken 2017).

The following procedure and set of assumptions is used to calculate carbon emission reductions associated with climate change education, as shown in Figure 5. We first assume that a modest investment in climate change education would allow students of secondary school age from middle and high income countries (where their carbon emissions are highest) to receive a specialized climate change education (i.e., using similar educational methodologies as we’ve ourlined here), and that students who receive this education would each reduce their carbon emissions by 2.86 tons of CO_2_/year (i.e., as in the COMM 168 class), for that year, and for each year following. Further, we assume that such a program would start small at 1 million students and grow by 13% per year until 2050, when the program reaches over 38 million participants. We use 2015 data from the United Nation Educational, Scientific and Cultural Organization (UNESCO) (http://data.uis.unesco.org) to estimate the number of students of secondary school age from high income and upper middle income as 298 million. This allows us to estimate the percentage of students participating in this specialized climate change education program in 2020 and 2050, assuming the population of secondary students in these countries does not change.

**Figure 5.**
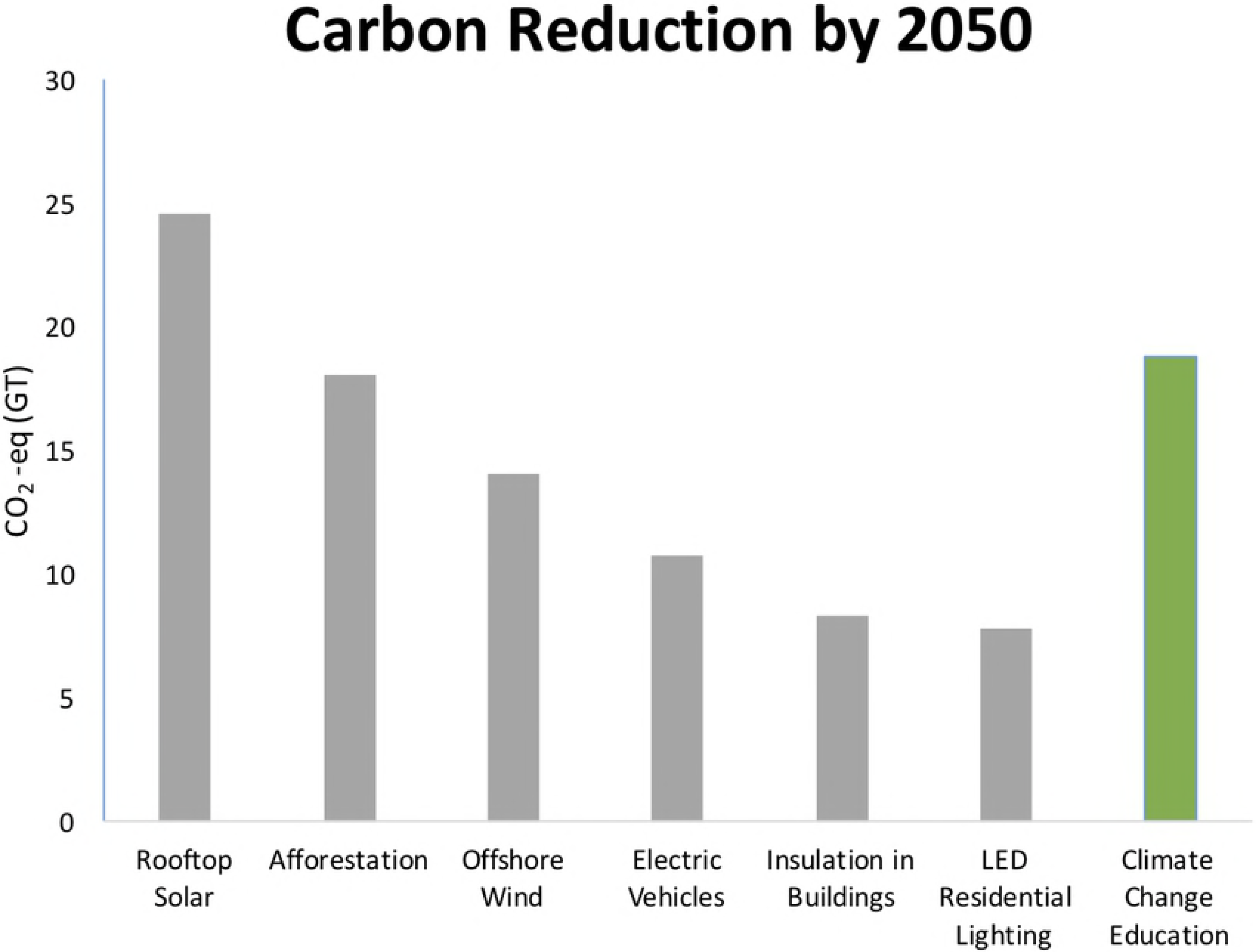
Comparison of various existing technologies that can be applied over a 30-year period (2020-2050) to help reduce global carbon emissions. The potential role of climate change education programs is calculated using the per student carbon reductions estimated from the COMM 168 class.

In Figure 5, six of the solutions presented in Project Drawdown are compared with our own estimate for using education as a climate change mitigation strategy. For the solution scenarios developed by Project Drawdown, each of these represents ambitious and yet, also technically and economically feasible plans for reducing carbon levels. Technical details and reference literature for all these solutions are presented at www.drawdown.org. As examples, the Rooftop Solar scenario grows the percentage of electricity generated by rooftop solar from 0.4% today to 7% by 2050, while the Electric Vehicles scenario grows the percentage of passenger miles from electric vehicles from less than 1% today to 16% by 2050. For the Climate Change Education scenario we assume that a) each student reduces their carbon emissions by 2.86 tons of CO_2_, similar to the COMM 168 class and b) the adoption of this type of education grows from less than 1% of all secondary students today to 16% of all secondary students by 2050 (note: the number of secondary students is restricted to only high income and upper-middle income countries where residents have higher carbon emissions.)

The results of this comparison show that education, if designed appropriately, can potentially be as effective as other established climate change mitigation techniques. Based on the scenario we developed, the implementation of climate change education over a 30-year period (2020-2050) could reduce emissions by 18.8 GT of CO_2_ eq, an amount that would rank in the top quarter (15 out of 80) of the presented solutions in Project Drawdown. Although at scale, the use of education as a climate change mitigation technique is still untested, our analysis suggests that if the educational approach is sound, and if we take the effort to measure the impact of education, we may realize the potential to reduce carbon emissions using education. We also acknowledge that although some barriers to developing a successful large-scale climate change education program exist, significant social and political challenges exist with most large-scale solutions to climate change.

## 5. Discussion: Application to K-12 educational environments

The above results describe the impact of an educational design on college students and the potential impact of scaling such a program across a large number of secondary school age students. We now discuss some initial work by the authors to develop K-12 curriculum that uses some of the design approaches identified in the COMM 168 class.

Within K-12 education, national and regional standards play an important role in guiding the design of classroom learning experiences. In the U.S., the National Research Council of the National Academy of Sciences helped develop an evidence-based framework for teaching science that included contemporary research on the ways students learn science most effectively (National Research Council 2012). As a result of the framework, the Next Generation Science Standards (NGSS) were released in 2013 as a state-led set of science standards that would be adopted at the state level. At present, 19 states have adopted the NGSS (representing 36% of students) and another 19 states have used the NRC Framework for K-12 Science Education in developing their own standards (representing 29% of students). These standards offer a new way of teaching science that emphasizes integrative learning across science fields and applies knowledge to real-world contexts. Given our experiences in the COMM 168 class and the guidelines presented in the NGSS, we saw an opportunity to teach the required science subjects (i.e., life science, physical science, earth science and engineering design) using climate change and environmental solutions as an overarching theme. Other research suggests that using environmental subjects can promote student learning and higher-order thinking (e.g., Ruiz-Gallardo et al. 2013; Dresner et al. 2014).

Over the last three years, the authors have been working to apply an educational model similar to that used in COMM 168 to a middle school science curriculum. In particular, the design of the curriculum emphasizes a) establishing a personal connection to students’ lives, b) providing students with opportunities to create change in their own communities and c) helping students develop empathy for nature and the environment. The results of this curriculum design include the creation of a student-relatable storyline (i.e., collection of videos and games featuring a climate action superhero) and the focus on learning experiences that help students build a personal connection to climate change through participation in experiences where students design climate solutions for their communities. Each unit of science instruction has a focus on solutions to an environmental problem, and a continuity of learning across a middle school sequence is designed to potentially provide students with a strong connection to environmental stewardship. The middle school curriculum is currently being used in a number of school districts in California and studies examining changes in student attitudes and behavior will be reported in the future.

As an example of the type of educational experiences being designed, we share some insights from an ongoing study to determine whether student motivation and agency around science and environmental solutions can be enhanced through a storytelling and filmmaking experience. During one of the five-week units of study, middle school students learn about the basics of climate science and are guided through the process of storytelling and filmmaking as a method of taking action on climate change. Students create their own short films about solutions to climate change, and these films are screened to friends, family and submitted to a regional film festival.

Overwhelmingly, students, teachers and parents reported that this unit of study was a valuable learning experience. Students enjoyed the filmmaking experience and the autonomy they had to create their own type of climate solution. This is reflected in student interviews indicating that their favorite part was that “we had no limitations at all” and “we got to put our storyboard and the ideas that we had into action.” Further, several students reported that they were either interested in the opportunity to do more community action, or that they were now taking on leadership roles towards further action. During an interview, one student responded, “Actually, um, after this project I’m, like, really considerate of what I’m using…what my family and I could do to, like, reduce global warming.” Given that most students reported that they did not know much about climate change prior to this unit, this feedback suggests that such measures applied in K-12 environments may be valuable tools for inspiring social change.

## 6. Conclusions

The potential role of education on individual carbon emissions was studied using five years of data from students who completed an intensive university course on climate change. Students were surveyed at least five years after having taken the course, and their responses were used to provide both qualitative and quantitative measures of the impact of the course on their attitudes and behavior regarding solutions to climate change. The university course was designed to be impactful, including various elements to engage students around personal and social activism. In open-ended feedback and the focus group interviews, students recounted how the class has changed their lives, both personally and professionally. Examples of personal changes included the type of car they drive and the type of food they eat. Examples of professional changes included how they create environmental benefits through their job. The results from the survey data also showed that the course was impactful, even many years later. Student behavior related to waste decisions, home energy decisions, transportation and food choices all showed significant behavior change that was attributed to the COMM 168 course, and these changes were quantified using a reputable online carbon emissions calculator. The carbon emissions for an average California resident are 25.1 tons/year, while the estimated reductions in carbon emissions from the COMM 168 graduates are 3.54 tons/year. It was found that the participants who had personally experienced the effects of global warming, or felt that global warming will harm them personally, had the largest reductions in carbon emissions.

This study suggests that the COMM 168 class provided at least two of the crucial factors that Hungerford and Volk (1990) identify as contributing to pro-environmental behavior: ownership variables and empowerment variables. Surveys and focus group interviews demonstrated that students in this course established in-depth knowledge of climate change and demonstrated knowledge of how to act in the service of climate change mitigation. Surveys and focus group interviews also revealed that graduates of the course feel a lasting personal connection to the issue and have confidence in the success of their actions. This strong sense of personal obligation and the perceived individual agency to address climate change suggests that education with an emphasis on community engagement may provide a public benefit.

The potential to use education as a climate change mitigation measure would be valuable and in line with other mitigation measures if such reductions as achieved in the COMM 168 course could be achieved in other classrooms. We demonstrate this through comparisons with other climate change solutions and show that at scale, climate change education can be as effective in reducing carbon emissions as other solutions such as rooftop solar or electric vehicles. The notion that education is an important part of responding to climate change is not novel, and yet rarely has it been quantified and measured. This paper sheds light on how such measurements could be taken, and it offers a pedagogical insight for how to make education an effective climate change mitigation strategy.

## Appendix A Survey instrument used for the graduates of the COMM 168 class

Intro: Thank you for agreeing to take our survey about the Global Climate Change Course (COMM/ENVS/GEOL/HUM/METR 168—hereafter COMM 168). This is divided into two sections: general information and post-course life.

General Questions:

1. What academic year did you enroll in COMM 168?

a. 2007-2008
b. 2008-2009
c. 2009-2010
d. 2010-2011
e. 2011-2012
2. What is/was your major?
3. When did you graduate from SJSU?

a. Spring 2008
b. Fall 2008
c. Spring 2009
d. Fall 2009
e. Spring 2010
f. Fall 2010
g. Spring 2011
h. Fall 2011
i. Spring 2012
j. Currently enrolled
4. How much do you think global warming will harm future generations?

a. Not at all
b. Only a little
c. A moderate amount
d. A great deal
e. Don’t know
5. How much do you think global warming will harm you personally?

a. Not at all
b. Only a little
c. A moderate amount
d. A great deal
e. Don’t know
6. Which of the following statements comes closest to your view?

a. Global warming is not happening.
b. Humans cannot reduce global warming, even if it is happening.
c. Humans could reduce global warming, but people are not willing to change their behavior, so we are not going to.
d. Humans could reduce global warming, but it is unclear at this point, whether we will do what is needed.
e. Humans can reduce global warming, and we are going to do so successfully.
7. Rate your opinion of the following statement: The actions of a single individual will not make any difference in global warming.

a. Strongly agree
b. Somewhat agree
c. Neutral
d. Somewhat disagree
e. Strongly disagree
8. Rate your opinion of the following statement: New technologies can solve global warming, without individuals having to make big changes in their lives.

a. Strongly agree
b. Somewhat agree
c. Neutral
d. Somewhat disagree
e. Strongly disagree
9. Which comes closer to your own view?

a. Most scientists think global warming is happening.
b. Most scientists think global warming is not happening.
c. There is a lot of disagreement among scientists about whether or not global warming is happening.
d. Don’t know enough to say.
10. How much do you agree or disagree with the following statement: “I have personally experienced the effects of global warming.”

a. Strongly agree
b. Somewhat agree
c. Neutral
d. Somewhat disagree
e. Strongly disagree
11. How many of your friends share your views on global warming?

a. None
b. A few
12. personally recommended COMM 168 to other students

a. True
b. False The following questions relate to your participation in COMM 168: Global Climate Change at San José State University, and if this course has affected any of your actions or decisions since you took the course.
13. As a result of my participation in this course, I have taken the following actions to reduce the amount of waste produced in my home. (Check all that apply):

□ Recycle more often
□ Buy products that have less packaging
□ Compost food scraps or food waste
□ Give away or donate products so that they can be reused
□ Other:
□ The course didn’t have an influence on these actions
14. As a result of my participation in this course, I have taken the following actions to reduce energy consumption at home (Check all that apply):

□ Purchased renewable energy from utilities
□ Purchased energy saving appliances
□ Installed solar P V
□ Installed solar hot water
□ Changed traditional light bulbs to energy-efficient light bulbs
□ Other:
□ The course had no influence on my home energy use
15. As a result of my participation in this course, over the course of a week, I make food choices to reduce carbon emissions

a. All the time
b. Often
c. Sometimes
d. Occasionally
e. The course had no influence on my food choices
16. As a result of my participation in this course, I have made the following changes to my transportation methods (Check all that apply):

□ Purchased a hybrid car
□ Carpool regularly
□ Purchased a more gas-efficient car
□ Used public transportation more often
□ Used a bicycle instead of a car as transportation method
□ The course didn’t have an influence on my transportation methods
17. As a result of my participation in this course, I purchase carbon offsets when I fly.

a. All the time
b. Often
c. Sometimes
d. Occasionally
e. Never
f. The course had no influence on my purchase of carbon offsets
18. 18. Was there any other situation when you remember how particular content of the class influenced a decision(s) in your life? Explain.

## Appendix B Estimating reductions in carbon emissions from the survey responses

The procedure used to estimate the impact of various behaviors on carbon emissions is described here. In the survey, participants described particular actions they have taken as a result of taking the COMM 168 class. We use an online carbon footprint calculator to estimate how those actions change an individual’s annual carbon emissions. The carbon footprint calculator we use was developed by the CoolClimate Network: a partnership between government, business, NGOs and universities including the University of California, Berkeley. This tool is supported and endorsed by the California Air Resources Board (Jones and Kammen 2011).

The CoolClimate Calculator is publicly available (http://coolclimate.berkeley.edu/calculator) and has been used in a number of studies (e.g., Shirley et al. 2012). The categories of questions to estimate the household carbon emissions include travel, home, food, goods and services. As shown in Figure B1, each of these categories includes subsections that include additional questions that allow for more accurate estimates of carbon emissions for each category. The calculator makes various assumptions based on user responses, but it also allows for more detailed responses through an advanced section. For example, in the housing section, the advanced section allows the user to enter detailed information about the building design (e.g., area and type of insulation including windows) and the type of appliances and heating/cooling systems, versus the regular analysis that estimates carbon emissions based on how much energy an average household uses.

**Figure B1.**
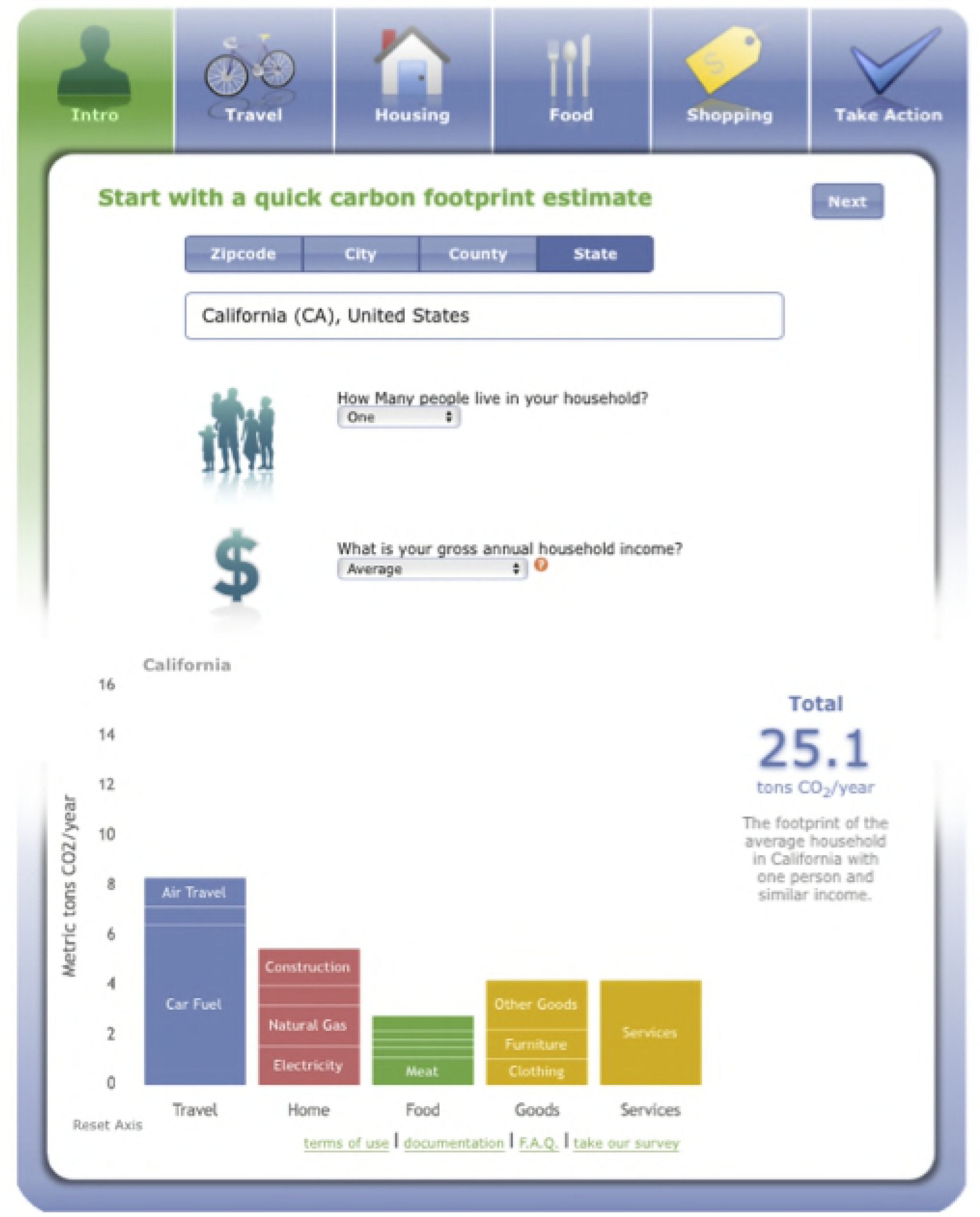
Screen shot from the CoolClimate calculator developed in part by the California Air Resources Board (www.coolcalifornia.org)

The following procedure is used to estimate the reductions in carbon emissions due to specific actions. We start by selecting one person with an average annual household income for the state of California. Given the internal assumptions based on the categories of travel, home, food, goods and services, the annual carbon emissions are 25.1 tons of CO2/year for the individual. We then select the ‘Take Action’ section, where changes in individual actions together with the underlying assumptions can be made for a number of categories. We use this section to match the responses from the COMM 168 participants with reductions of carbon emissions, using a set of assumptions that are shown in Table A1. Each assumption comes from the CoolClimate calculator and provides an estimate of how a particular action would reduce annual carbon emissions. For example, one of the survey questions asks if the participant purchased renewable energy from their utility. The CoolClimate calculator assumes the household uses 4,396 kWh of electricity/year, which produces 1.36 tons of CO_2_. We assume that participants who answer ‘yes’ to the question about purchasing renewable energy from their utility would get 100% of their electricity from clean energy and thus would save 1.36 tons of CO_2_/year. In another example, for the question about purchasing a more efficient vehicle, the calculator assumes that the individual would trade in a 22 mpg vehicle for a 32 mpg vehicle, and after driving 13,100 miles/year, they would save 2.08 tons of CO_2_/year. Each of the question responses and associated assumptions are given in Table B1.

**Table B1.** Description of the assumptions and reductions in carbon emissions for each of the survey responses.

| **Question Response** | **CO<sub>2</sub> reduction (tons/year)** | **Assumptions used in CoolClimate calculator** |
| --- | --- | --- |
| Buy renewable energy | 1.36 | 100% electricity from renewables |
| Buy energy star appliances | 0.04 | Energy star fridge or other products |
| Install solar panels | 0.68 | 50% electricity from renewables |
| Install solar hot water | 0.4 | 50% of heating of water from solar hot water |
| Change light bulbs | 0.16 | 5 bulbs used 5 hours per day |
| Buy hybrid/electric car | 4.22 | average of hybrids (40 mpg*2/3) + electric vehicles (99 mpg*1/3)) = 59.67 mpg |
| Recycle more often, buy products with less packaging, compost food scraps, give away/donate products | 0.42 | Reduce waste by 25% for each action |
| Carpool regularly | 0.85 | 3 times/week |
| Buy more fuel efficient vehicle | 2.08 | 32 mpg vs. 22 mpg for 13,100 miles/yr. |
| Use public transit more | 0.42 | 20 miles/week in bus instead of 22 mpg in car |
| Use bike for transportation | 0.53 | 20 miles/week instead of 22 mpg in car |
| Make food choices to reduce emissions | 0.69 | Response of 'all the time' or 'often' is default setting for 'low carbon version of American diet' (0.69 tons/year). Response of 'sometimes' or 'occasionally' is a reduction of 0.46 tons/year through increase in meat consumption from 244 calories (default) to 353 calories. |
| Buy carbon offsets for flying | 0.93 | Response of 'all the time' or 'often' is 80% of flights purchased offsets (0.93 tons/yr.). Response of 'sometimes' or 'occasionally' is 40% of flights purchased offsets (0.46 tons/yr.). |

## Acknowledgements

The authors would like to thank Dr. Elizabeth Walsh for her helpful suggestions on our data analysis and the National Science Foundation (Grant #1513332) for supporting our K-12 science education research. Dr. Cordero acknowledges that he is the majority owner of Green Ninja Inc.

